# Electrodeposited Biocompatible Coatings for 3D Electrodes on Retinal Prostheses

**DOI:** 10.64898/2026.09.06.749620

**Authors:** Viktoryia Shautsova, Emma Butt, Mohajeet Bhuckory, Linh Mai Vu, Davis Pham-Howard, Vladimir Mamchik, Ludwig Galambos, Jeonghyun An, Andrew Shin, Keith Mathieson, Theodore Kamins, Daniel Palanker

## Abstract

Photovoltaic subretinal prosthesis, PRIMA, provides central vision to patients blinded by age-related macular degeneration, with acuity matching the 100 μm pixel size. Further miniaturization requires pillar electrodes to position the stimulating surfaces closer to the inner retinal neurons. While such structures can be electroplated in gold and coated on their tops with SIROF, the exposed gold sidewalls are not biocompatible. Sputtering or atomic layer deposition of protective coatings are unsuitable for selectively passivating the pillar structures without also coating the photosensitive regions and return electrodes of the implant.

Here, we present a strategy for biocompatible coating of pillar sidewalls while preserving surrounding implant functionality. The approach combines non-critical photoresist lithography to protect planar return electrodes with electrodeposition of TiO₂ or Pt onto gold pillar sidewalls. In-vivo studies demonstrated that both TiO₂ and Pt coatings are biocompatible and prevent adverse reactions of the retinal tissue to gold. Since specific capacitance of electroplated TiO₂ (25 μF/cm²) is much lower than that of Pt (240 μF/cm^2^), the former better limits the current from the side walls and ensures that charge injection occurs predominantly through the pillar tops coated with SIROF (∼6 mF/cm²).

Electrodeposition, combined with noncritical photolithography provides a scalable wafer-level solution for fabrication of biocompatible three-dimensional electro-neural interfaces, addressing a critical bottleneck in bioelectronics.

## 1. INTRODUCTION

Patients afflicted by retinal degenerative diseases, such as age-related macular degeneration (AMD) and retinitis pigmentosa (RP), experience loss of vision due to death of photoreceptors, while inner retinal neurons are preserved to a large extent [1, 2]. Retinal prostheses aim at restoring visual function by electrical stimulation of the remaining inner retinal neurons. The retinal prothesis system PRIMA demonstrated restoration of central vision in patients blinded by atrophic age-related macular degeneration with resolution matching its 100gm pixel size (20/420) [3]. The system includes a wireless photovoltaic subretinal implant and a pair of augmented reality (AR) glasses equipped with a video camera (Figure 1a). Images captured by the camera are processed and projected onto the implant using intensified pulsed light. To avoid stimulation of the healthy peripheral retina, the glasses use an invisible near-infrared (NIR, 880nm) wavelength. The 30 gm thick silicon implant is composed of hexagonal photovoltaic pixels that convert pulsed light into electric current to stimulate the nearby neurons in the inner nuclear layer (INL), primarily the bipolar cells (BCs). The 2x2 mm arrays are intended for clinical use, whereas a 1.5×1.5 mm version is used for process development and studies in rats.

**Figure 1.**
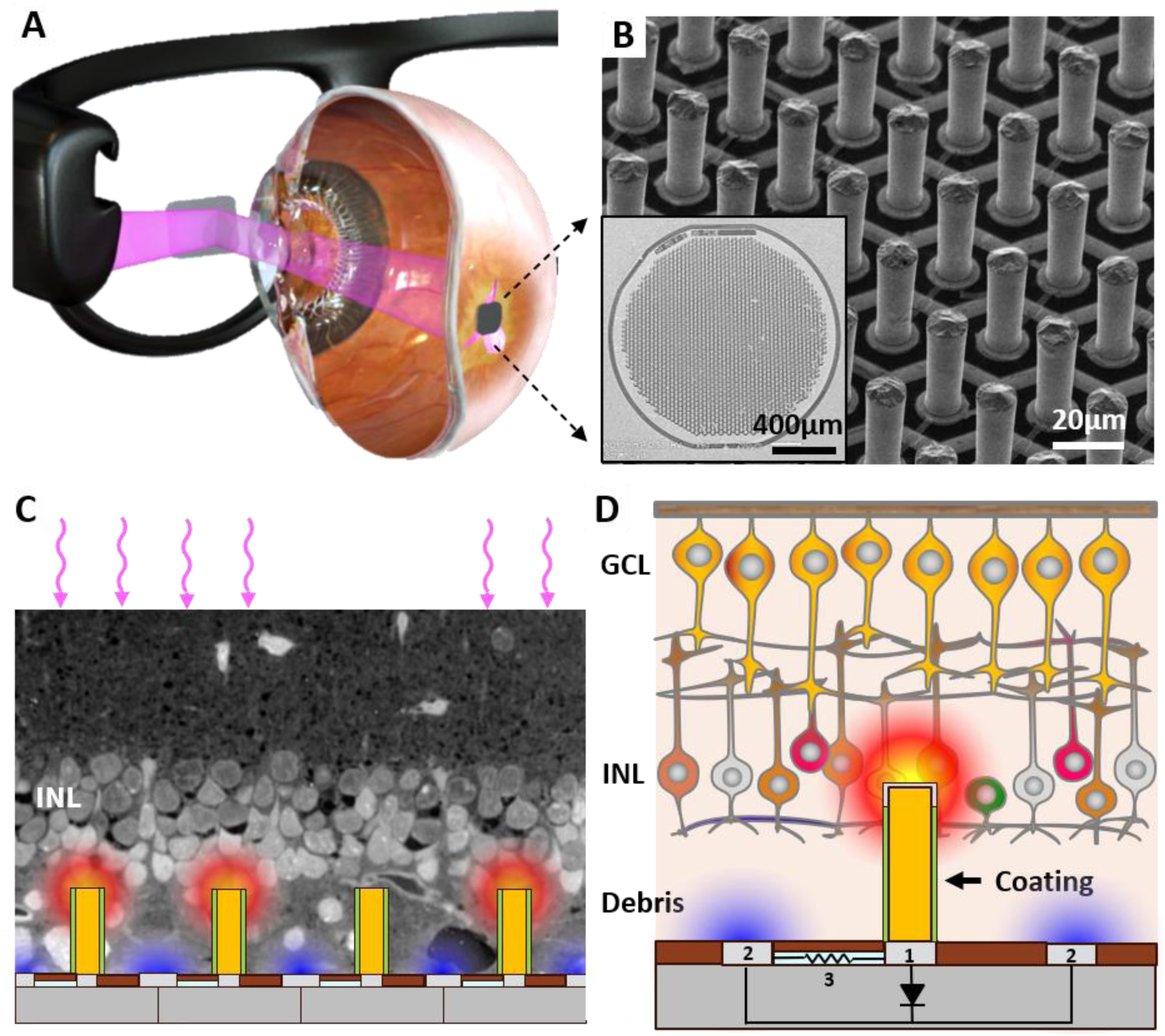
System diagram. A) Images captured by video camera are processed and projected onto subretinal implant using pulsed near-infrared (880 nm) light. Photovoltaic pixels convert it into electrical current to stimulate the bipolar cells in the Inner Nuclear Layer (INL). B) SEM images of the typical fabricated implant (inset) with pillar electrodes providing better proximity to INL. C) Illustration of electric field generated by the current flowing from the top of pillars to return electrodes at the base, overlayed with histology of a degenerated retina. The exact potential distribution in electrolyte calculated using COMSOL Multiphysics is provided in Supplementary Note S1. D) Pillars are deposited on top of the active electrodes (1) and current is collected by return electrodes (2), connected in a mesh. To discharge the electrodes between pulses, active electrodes are connected to return electrodes via shunt resistor 3. Electroplated pillars can penetrate the debris layer to deliver current to bipolar cells in the INL. To avoid direct contact between the pillar sidewalls and tissue, they require a low-capacitance biocompatible coating.

Despite being the highest prosthetic acuity to date, further improvements in resolution are needed to help a larger population of AMD patients since their remaining peripheral vision often supports visual acuity of no worse than 20/400. For acuity exceeding 20/100, pixels below 25 µm would be required. In rats, arrays with pixels down to 20 µm demonstrated a grating acuity up to their natural resolution limit of 27 µm [4]. However, unlike rats, the human degenerate retina has a debris layer of 20-60 µm in thickness, which separates the INL from the subretinal implant [5, 6]. This separation increases the stimulation threshold and decreases the contrast and resolution.

Pillar electrodes offer a solution to this problem, as tissue migration into the voids between the pillars of the subretinal implant enables closer proximity of electrodes located on top of the pillars to the target neurons (Figure 1b) [7, 8]. Previously, we introduced a wafer-scale, cost-effective technique for fabricating 40gm tall electrodes by gold electroplating on top of the photovoltaic arrays [9]. To improve biocompatibility, we also investigated electroplated platinum pillars, which suffered from stress-induced failure, including cracking, peeling and bending [10]. For these approaches, the top of each pillar is coated with high-capacitance sputtered iridium oxide film, SIROF, enabling the charge injection primarily from the top of the 3-D electrode, close to the target neurons [9], obviating the need for insulating the side walls.

While gold offers excellent conductivity and compatibility of the pillars’ fabrication with other processes in manufacturing of photovoltaic arrays, the exposed side walls of gold pillars present several problems. First, retinal tissue exhibits adverse reaction to gold even with no electrical current [11]. Second, electrochemical reactions on gold electrodes can lead to their erosion and tissue toxicity, particularly under chronic stimulation [12, 13]. Therefore, a protective biocompatible coating is required to isolate the structural metal from the surrounding tissue (Figure 1c,d). Such coating should have low capacitance to confine the current injection to the SIROF-coated pillar tops and suppress the undesired charge injection from the sidewalls [9]. The biocompatible coating should exhibit long-term stability *in-vivo* and be suitable for high aspect ratio 3D fabrication on a wafer scale.

Deposition of a conformal coating on 3D structures is challenging for conventional microfabrication techniques, especially if it can harm the active structures on the device, such as photosensitive regions and return electrodes in our photovoltaic implants. Defining photolithographic lift-off layers around tall, narrow 3D structures is particularly problematic, as a nonuniform resist coverage compromises the pattern transfer during sputtering, often leading to incomplete coverage near the pillars, resulting in unintended coating of photosensitive areas. While atomic layer deposition (ALD) provides excellent conformity on high aspect ratio structures, its practical integration into complex device architectures is still constrained by difficulties with lift-off processes [14] and a lack of selective deposition [15], often resulting in inadvertent insulation of electrically active regions, such as return electrodes in a retinal prosthesis. Together, these limitations highlight the need for novel fabrication strategies of high aspect ratio electrodes that combine deposition conformity and spatial selectivity, while being cost-effective, high-yield and compatible with wafer-scale processing.

In this work, we present strategies for wafer-scale, cost-effective fabrication of biocompatible coatings on pillar electrodes leveraging electrodeposition techniques to selectively cover electrically addressable surfaces. This selectivity allows the use of noncritical photolithography masks, substantially relaxing the alignment and patterning constraints associated with high– aspect-ratio geometries. Selective electrodeposition of TiO_2_ and Pt on gold pillar electrodes provides a stable and biocompatible coating, allows full integration with the existing photovoltaic retinal implant design, and is expected to enable scaling pixel sizes from 100 to 25 gm in human patients. More broadly, fabrication strategies presented here are applicable to a wide range of complex electro-neural interfaces and provide a scalable pathway to engineering stable, low-capacitance protective coatings for 3-D neural prostheses.

## 2. RESULTS AND DISCUSSION

### 2.1. Selective coating of side walls on high-aspect-ratio electrodes

Gold pillar electrodes of 40 gm in height with diameters of 22, 16.5, 11.5 and 8.5 µm were electroplated on active electrodes in hexagonal pixels of 55, 40, 30 and 22 µm in width, respectively, as described previously [9,15] (Figure 2A,D). Each array was 1.5 mm in diameter and return electrodes were interconnected across the entire wafer, providing electrical access for electroplating of the pillars and subsequent electrodeposition of the coatings via shunt resistors connecting the central disk electrode in each pixel to the return electrode mesh (Figures 1D, 2A). Pillar electroplating was performed using a template made from a 35 gm thick negative photoresist layer, PR1, (Figure 2A,D), as described in our prior work [9].

**Figure 2.**
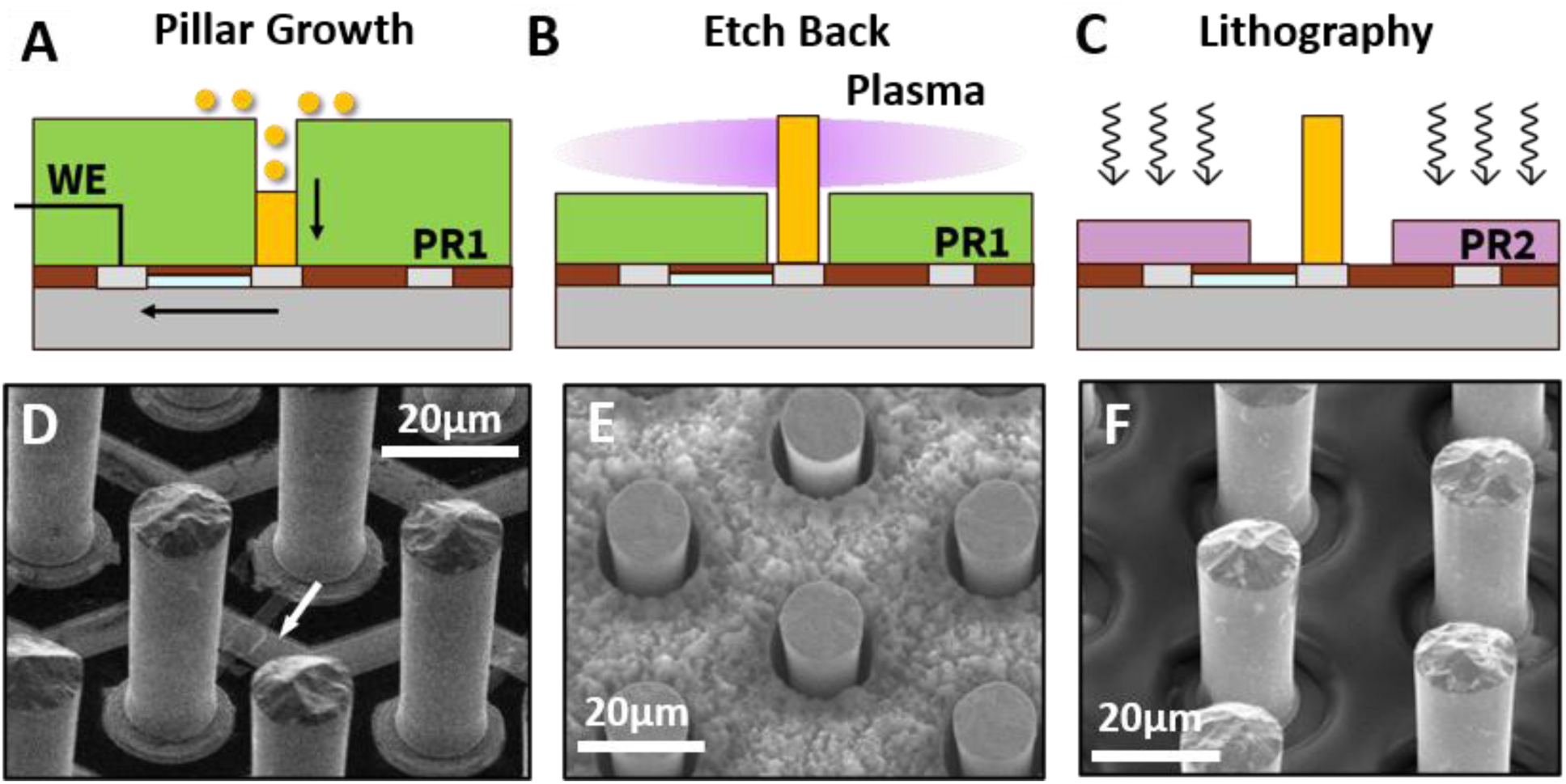
Pillar electroplating and fabrication of noncritical photoresist masks for electrodeposition. A) Pillar electroplating is performed using a photoresist template (PR1) patterned to expose the active electrodes. Electric current flows via the mesh of return electrodes and shunt resistors to the electroplating sites. B) Etch-back: The same photoresist layer (PR1) is etched back to expose the sidewalls of the gold electrodes. C) Lithography: after stripping PR1, a second photoresist layer (PR2) is patterned using noncritical photolithography to open windows around the pillars, while protecting the underlying return electrodes. D-F) SEM images of representative devices with 40µm-high gold pillars, shown after PR1 removal (D), after etch-back process (E) and after noncritical photolithography process (F).

For selective electrodeposition of the protective coating on pillar sidewalls, the return electrodes should be protected, while the sidewalls remain fully accessible to the solution. For this purpose, two alternative fabrication strategies were explored, both producing noncritical masks where the photoresist edge is deliberately positioned away from the pillars, partially exposing the SiC-coated substrate. In the first approach, pillar sidewalls were exposed by removing the original PR1 photoresist around the pillars using a reactive ion etching process, RIE etch-back, over entire wafer (Figure 2 B,E). The photoresist etch was more pronounced near the pillars, exposing their surface all the way to the bottom, likely due to a local acceleration of the etch rate at the base of a conductive feature due to the concentration of deflected ions, called the microtrenching effect [16, 17]. The remaining photoresist above the rest of the device protects the return electrodes.

In the second approach, PR1 was completely removed after the pillar formation and replaced with a second photoresist layer (PR2). The negative photoresist layer, PR2, was patterned using a maskless direct-write photolithography technique (see Experimental Section for details) to form a noncritical mask that similarly exposed the conductive pillars and some insulated areas around them, while protecting the return electrodes (Figure 2 C,F). It is important to note that use of a positive photoresist for this purpose would require exposure adjacent to the pillars, which would lead to significant light scattering, resulting in poor pattern fidelity.

Both patterning approaches provide access to the pillar sidewalls for subsequent electrodeposition of biocompatible coatings, while protecting the return electrodes. Importantly, electrodeposition confines the material growth to exposed conductive regions, while avoiding deposition on insulated areas, such as exposed SiC above the photosensitive areas of the device. TiO_2_ and Pt were selected due to their widespread use in biomedical implants, established electrodeposition processes and well-known biocompatibility [10, 12].

### 2.2. Deposition and characterization of TiO₂ coatings

Titanium dioxide (TiO_2_) is widely used as an insulating coating in biomedical implants [18, 19]. As a wide-bandgap semiconductor (∼3.0–3.2 eV), TiO_2_ is expected to reduce current from the pillar sides and provide biocompatible interface with the retina. To form a thin TiO_2_ coating on Au, we adopted a well-established electrodeposition method based on anodic hydrolysis of a 50 mM TiCl_3_ solution [20]. The process was performed using a three-electrode setup under nitrogen and at pH of ∼2.3. As previously reported, the solution pH strongly influences the film morphology: moderate acidic pH values around 2.5 promote smooth films, while highly acidic solutions, such as pH ∼1, typically result in out-of-plane structural features, such as dendritic growth [21]. In our device, a smooth and uniform TiO_2_ surface is essential to preserve the intrinsically low capacitance of the coating and ensure that the current injection is predominantly from the SIROF-coated pillar top [9].

To study the properties of the TiO_2_ film grown on smooth Au substrates, we started with large planar samples, where deposition is well controlled. Using photolithography, a 5 mm diameter circular region on a gold substrate was exposed to electrodeposition solution. SEM imaging revealed a clear contrast between the bare Au and the deposited TiO_2_ film, which forms a continuous, crack-free coating (Figure 3A,B). Step-height analysis by AFM shows that the coating reaches ∼30 nm in thickness within the first 5 min of deposition, followed by a steady growth to ∼180 nm after 30 min and saturating around ∼200 nm at 60 min, indicating a self-terminating process (Figure 3C). Topography maps (Figure 3D,E) reveal that nanoscale features of the Au surface are substantially modified by electrodeposition, consistent with the formation of a continuous oxide layer. Correspondingly, the RMS roughness increases from ∼1.5 nm on Au substrate to ∼4 nm on TiO_2_ film, which remains largely constant despite the increasing film thickness (Figure 3F). Elemental analysis corroborates the formation of TiO_2_on the Au substrate. Energy-dispersive X-ray spectroscopy, EDS, confirmed the presence of Ti and O in the deposited film, as evident from the increased intensity of the Ti and O peaks (Figure 3G and Supplementary Note S2). X-ray photoelectron spectroscopy, XPS, shows the expected Ti 2p and O 1s signatures corresponding to stoichiometric TiO_2_ following removal of the surface layer by sputtering (Figure 3H and Supplementary Note S2).

**Figure 3.**
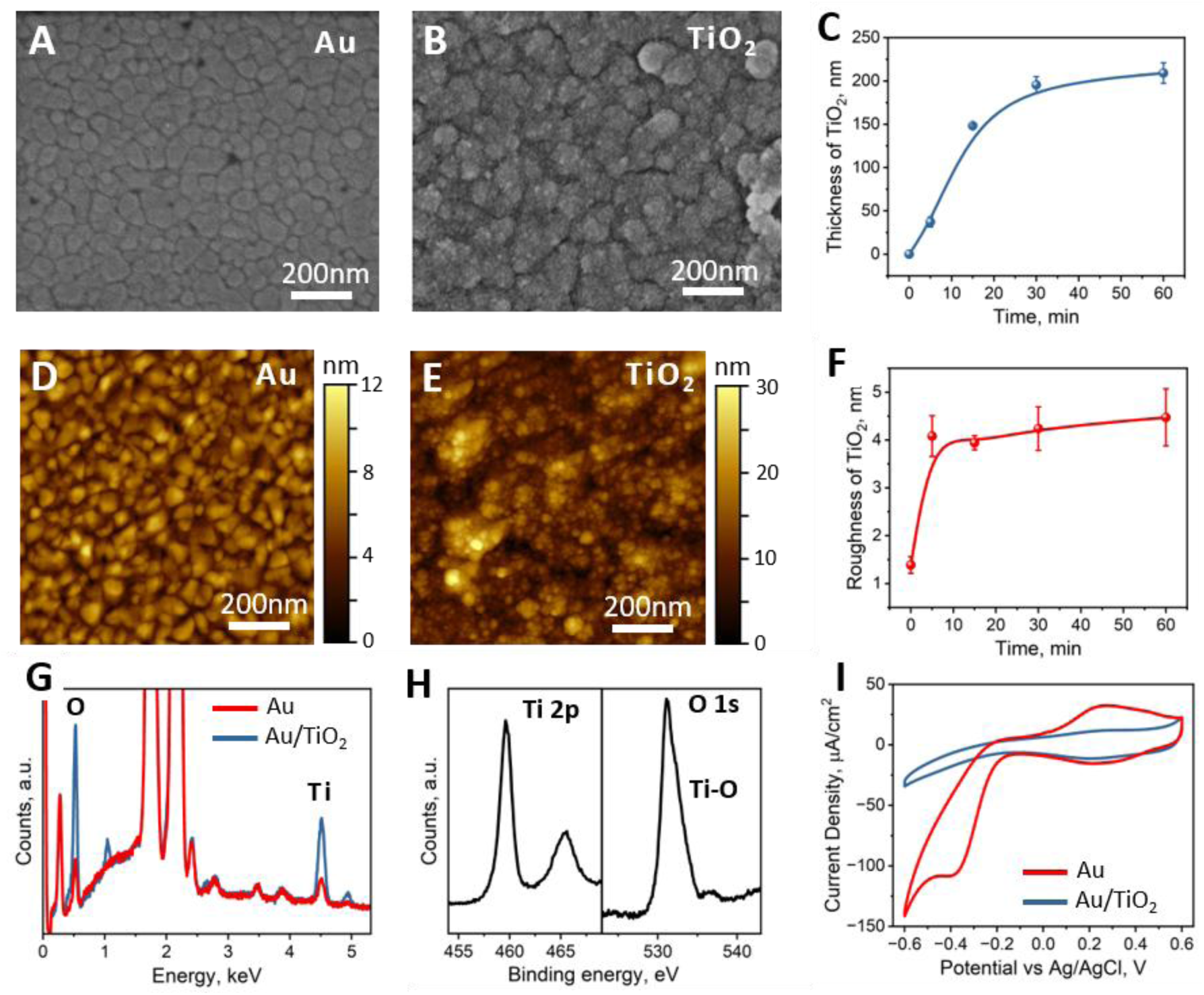
Properties of the electrodeposited TiO_2_ film. A-B) SEM of the Au substrate and the electrodeposited TiO_2_ film. C) Thickness of the TiO_2_ layer as a function of the deposition time, extracted from AFM step-height measurements. D-E) Surface texture of the Au substrate and the TiO_2_ film mapped by AFM topography. F) Surface roughness (RMS value), as a function of the deposition time, obtained from AFM analysis. G) EDS spectra verifying the presence of TiO_2_ film on Au substrate. H) XPS spectrum confirming the elemental composition consistent with TiO_2_, including the characteristic Ti 2p and O 1s features. I) Cyclic voltammetry comparing bare Au with electrodeposited TiO_2_film.

Electrochemical characterization demonstrates a decrease in charge injection from TiO_2_ compared to Au electrodes: Cyclic voltammetry (CV) shows suppression of the current injection, especially at negative bias, compared to Au (Figure 3i), with the same trend observed across all investigated electrodeposition times (Supplementary Note S3). Specific capacitance of the TiO_2_ film extracted from the CV data is approximately 25 μF/cm² (see Experimental Section), which aligns with previously reported values for amorphous TiO_2_ films and reflects the low-leakage interface desirable for electrode insulation [22, 23].

After establishing electrodeposition parameters for smooth TiO_2_ coatings on planar Au substrates, the process was applied to 3D pillars, using a noncritical photolithography mask as discussed above (Figure 4A). Prior to electrodeposition, an additional O_2_ plasma treatment was performed to remove the residual photoresist and prepare the pillar surface (Figure 4B). The TiO_2_ coating was then grown using the same electrical circuit as in the pillar electroplating.

**Figure 4.**
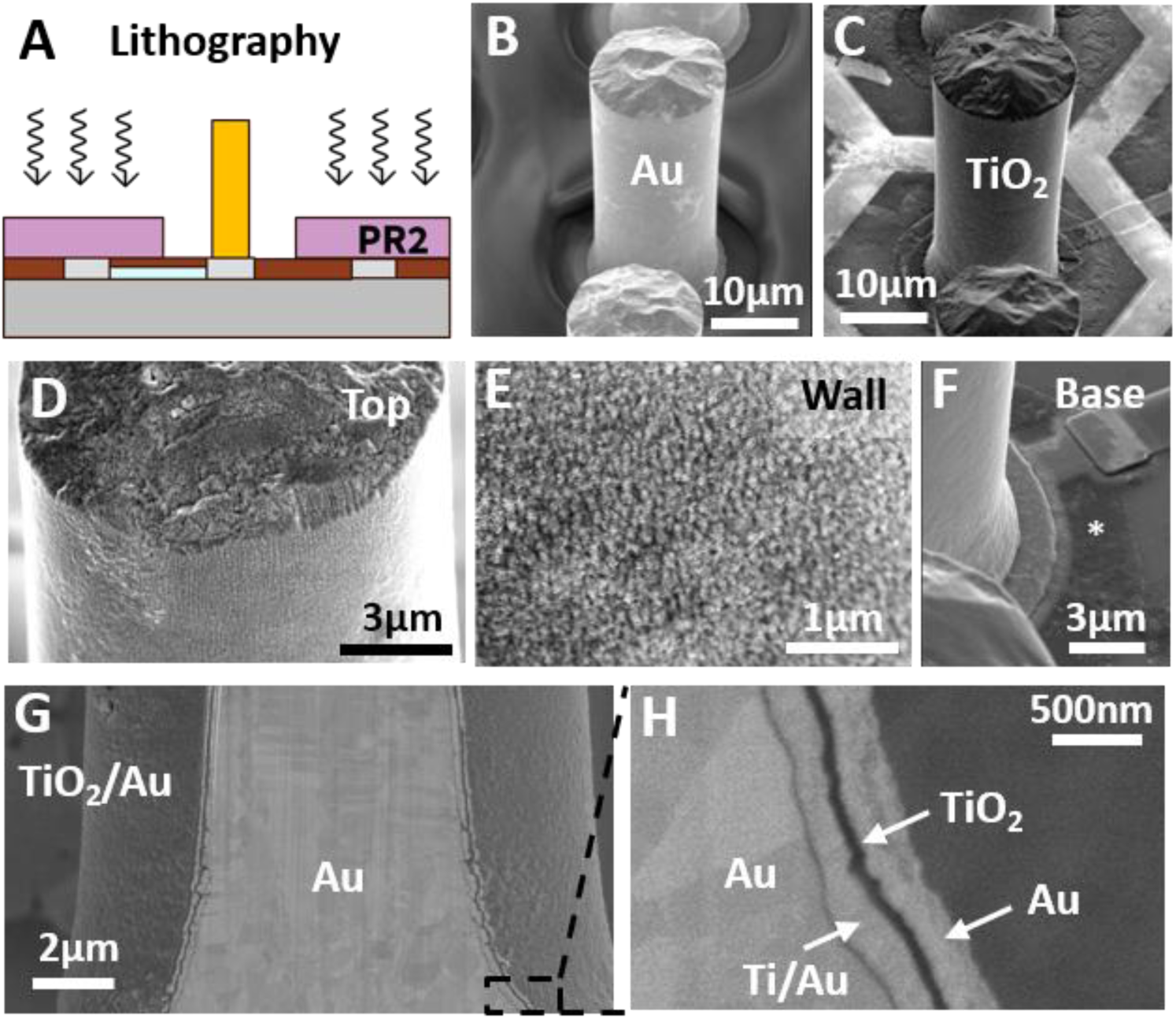
Electrodeposited TiO_2_ coating on pillar electrodes. A) Schematic of the noncritical photolithography approach. B) SEM demonstrates exposed pillars and negative photoresist (PR2) protecting the return electrodes. C-E) Pillar electrode after TiO_2_ electrodeposition. F) A thin TiO_2_ layer deposited from the solution onto insulated area around the pillar (*). G) Focused ion beam (FIB) cross-section of a pillar, revealing layers of electrodeposited TiO_2_, protected with sputtered Au for milling. H) The zoom-in image shows a continuous TiO_2_ layer, of about 30 nm in thickness. Note that the additional Ti/Au layer on top of electroplated pillars was used only for FIB to ensure an electrical connection for TiO_2_ electroplating.

SEM imaging demonstrated a uniform and continuous TiO_2_ coating along the pillar surface with no observable cracking or delamination (Figure 4 C-E). Similarly to planar substrates, EDS analyses showed the presence of the Ti and O peaks in the elemental composition of the deposited film. Focused ion beam (FIB) cross-sectional analysis further confirmed that the TiO_2_ electrodeposition results in a conformal coating around the Au pillar with a thickness of approximately 30 nm (Figure 4 G-H), consistent with observations on flat Au substrates, discussed earlier. A thin TiO_2_ layer was also detected on the SiC insulated flat base of the device (Figure 4f, indicated by *), attributed to passive deposition from the solution. Reflectance measurements demonstrated that this layer has negligible influence on optical reflectance, likely due to its minimal thickness (Supplementary Note S4).

Collectively, these results demonstrate that electrodeposition produces a continuous smooth TiO_2_ coating on planar substrates and 3D pillars, suitable for subsequent integration into the device manufacturing process.

### 2.3. Deposition and characterization of Pt coatings

Platinum (Pt) is one of the most established materials for implantable electrodes, owing to its excellent biocompatibility, chemical stability and efficient charge injection [10, 12, 24]. However, the high capacitance of Pt (around 0.1 mC/cm^2^ [25]) may reduce the confinement of the current flow around the SIROF-coated tops of the pillars [9]. Furthermore, electrodeposited Pt can result in rough surface topologies that serves to increase the capacitance of the interface and motivates experiments to check whether such a coating is suitable for 40gm tall pillar electrodes.

First, electrodeposition parameters were evaluated in Pt films grown on smooth planar Au substrates. A 5 mm diameter circular region was photolithographically patterned on a gold-coated substrate, and Pt was electrodeposited in chloroplatinic acid (see Experimental Section for details). The resulting films (Pt-ED) were compared to sputtered Pt, commonly used in neural electronics, as shown in Figure 5. Step-height analysis shows that ∼35 nm thickness was achieved after 20 minutes of growth at 1 mA/cm^2^, while lower current densities did not produce measurable growth under these conditions. Unlike TiO_2_, electrodeposition of Pt is not self-terminating and, therefore, the film thickness continues to increase with plating time. SEM (Figure 5 A,B) and AFM (Figures 5 D,E) of the platinum films reveal that Pt-ED coatings have a slightly rough surface populated by spherical structures, reaching 70 nm in diameter. RMS roughness of Pt-ED was about 7 nm, as opposed to 2 nm in sputtered Pt (Figure 5C). Pulsed Pt electrodeposition, previously reported as a route to smoother films [26], was also evaluated. However, AFM measurements showed no reduction in surface roughness (Supplementary Note S5).

**Figure 5.**
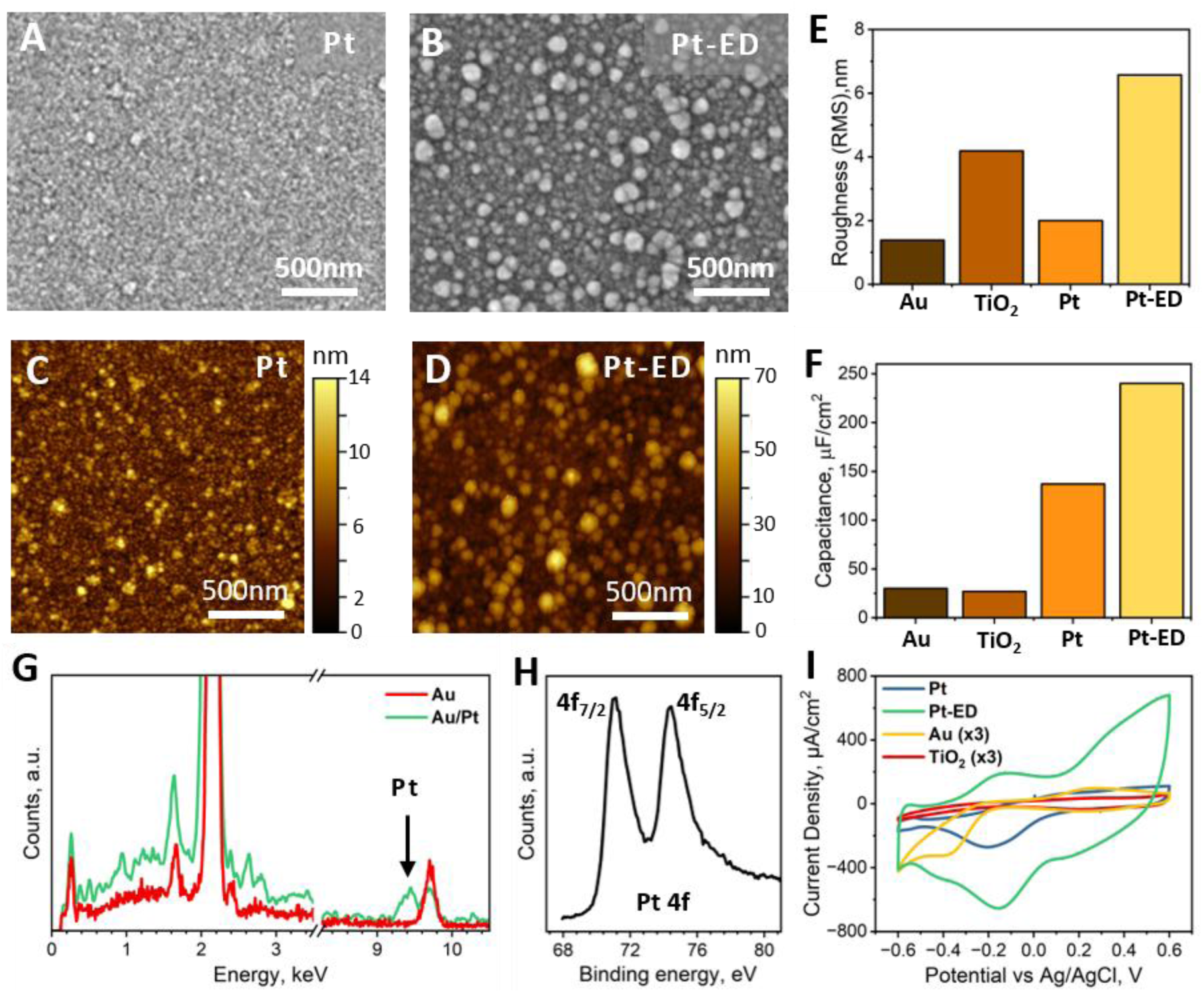
Electrodeposited Pt coating. A-B) SEM images of sputtered and electrodeposited Pt, respectively. C-D) AFM analysis of the surface texture mapped on sputtered and electrodeposited Pt, respectively. E) RMS roughness of the evaporated Au, sputtered Pt, electrodeposited Pt and TiO_2_ films. G-H) EDS and XPS measurements of electrodeposited Pt on Au. I) Cyclic voltammetry of evaporated Au, sputtered Pt and electrodeposited films of Pt and TiO_2_, and the corresponding specific capacitance (F).

XPS analysis of Pt coating revealed the characteristic 4f peaks at 71 (4f_7/2_) and 74.9 (4f_5/2_) eV, confirming that electrodeposited coating is composed of metallic Pt with a thin oxide layer on the surface, as confirmed by depth-profiling XPS (Figure 5H), and with no detectable residual chlorine or other byproducts from the electrodeposition solution (Supplementary Note S6). Since XPS typically averages over a relatively large area (∼100gm), EDS measurements were also performed on planar samples to establish a microscale elemental fingerprint of the Pt coating for subsequent analysis of coated pillars. As shown in Figure 5G, the uncoated gold typically exhibits only the peak at Lo=9.7 keV, while Pt-coated gold has an additional peak at Lo=9.4 keV - a characteristic EDS signature of Pt. Cyclic voltammetry (Figures 5I) reveals a specific capacitance of 240 µF/cm^2^ for Pt-ED films (Figure 5F), twice higher than specific capacitance of sputtered Pt, likely due to the increased roughness of the electrodeposited film (Figure 5D vs. 5C).

To perform Pt deposition on gold pillars, noncritical masks were first defined using two approaches described above: photoresist etch-back process (Figure 6A-C) and noncritical photolithography (Figure 6D-F). The etch-back step was carried out for 15 minutes to achieve uniform exposure of all gold pillars across the entire 4-inch wafer of devices. For the noncritical photolithography process, the conditions were identical to those used for TiO_2_ deposition discussed in the previous section. In both cases, a short O_2_ plasma clean, followed by an IPA rinse, was performed to clear any photoresist residue prior to electrodeposition.

**Figure 6.**
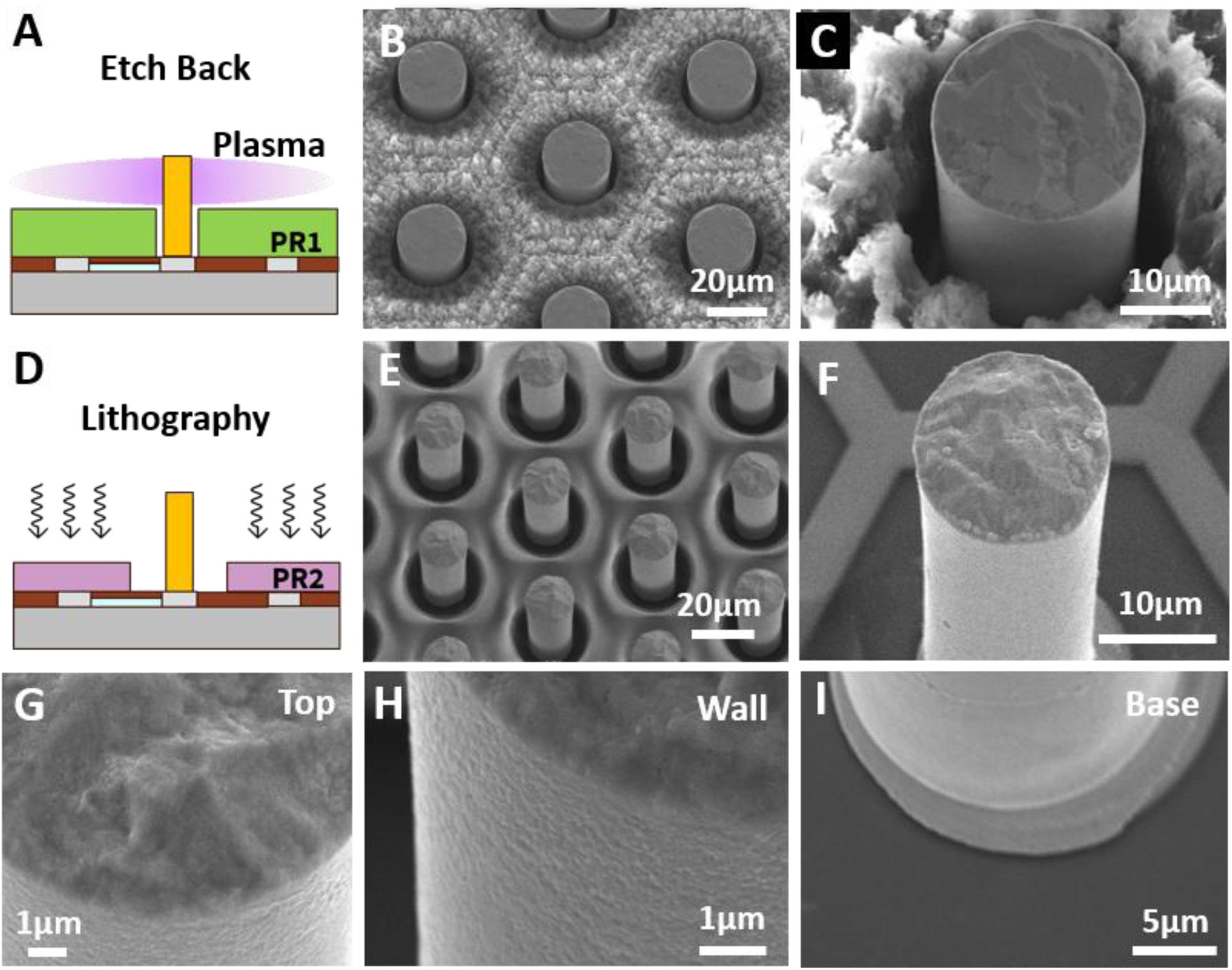
Electrodeposition of Pt on pillar electrodes using both methods for patterning noncritical photoresist masks. A) Schematic of the etch-back approach. B) Pillars and photoresist after etch-back process and O_2_/IPA cleaning. The pillars are exposed down to their base, whilst protecting the underlying return electrodes. C) Pillar electrode fully coated in Pt film before removing photoresist. D) Schematic of the noncritical photolithography approach. E) Au pillars and PR2 protecting the return electrodes prior to electrodeposition of Pt. F) Pt-coated pillar electrode after photoresist removal. G-I) Higher magnification SEM images demonstrating the continuous and smooth coating over the entire pillar height with no noticeable staining of the SiC at the pillar base.

Platinum electrodeposition was then carried out on the pillar electrodes, followed by photoresist removal. Using the Pt signature established above, correlative SEM-EDS measurements were performed at multiple positions along the pillar to evaluate continuity of the Pt coating, with particular attention to the pillar base, where access of the electrodeposition bath to the Au surface is most restricted. These measurements, corroborated by AFM results of continuous Pt-ED films on planar substrates, collectively suggest the Pt coating forms in a continuous manner over the pillar walls.

Both patterning techniques yielded uniform Pt coating across the entire pillar, extending down to the pillar base as confirmed by EDS (Supplementary Note S7). This is particularly critical for the etch-back process, where the surrounding photoresist is closer to the pillar and could potentially restrict the solution access during electrodeposition.

Higher magnification SEM showed similar roughness of the Pt coating to that of the planar Pt-ED films (Figure 6G,H), suggesting that capacitance of the formed Pt-ED coating is expected to be similar as well. Unlike the TiO_2_ electrodeposition process, no deposits were observed on SiC after platinum electrodeposition.

These results demonstrate the feasibility of platinum electrodeposition on high-aspect ratio gold structures. Importantly, both patterning techniques, the noncritical lithography method and the etch-back process, yielded continuous Pt coatings, offering flexibility in processes for integration with the rest of the device manufacturing flow.

### 2.4. Biocompatibility of Au, Pt and TiO_2_ coatings

Network-mediated stimulation by subretinal photovoltaic prostheses relies on intact bipolar cells as the primary targets, thus preservation of INL cells and their health is essential for functional efficacy. To determine the effects of pillar coating material on inner retinal cells, we examined the number of preserved cells, their morphology and microglial responses in retinal wholemounts 4-6 weeks after implantation in rats (Figure 7). Explanted retinas were immunohistochemically stained with fluorescent markers: PKCα for rod bipolar cells and IBA1 for microglia. We also used DAPI nuclear staining to assess the number of cells and TUNNEL staining to evaluate the extent of apoptosis.

**Figure 7.**
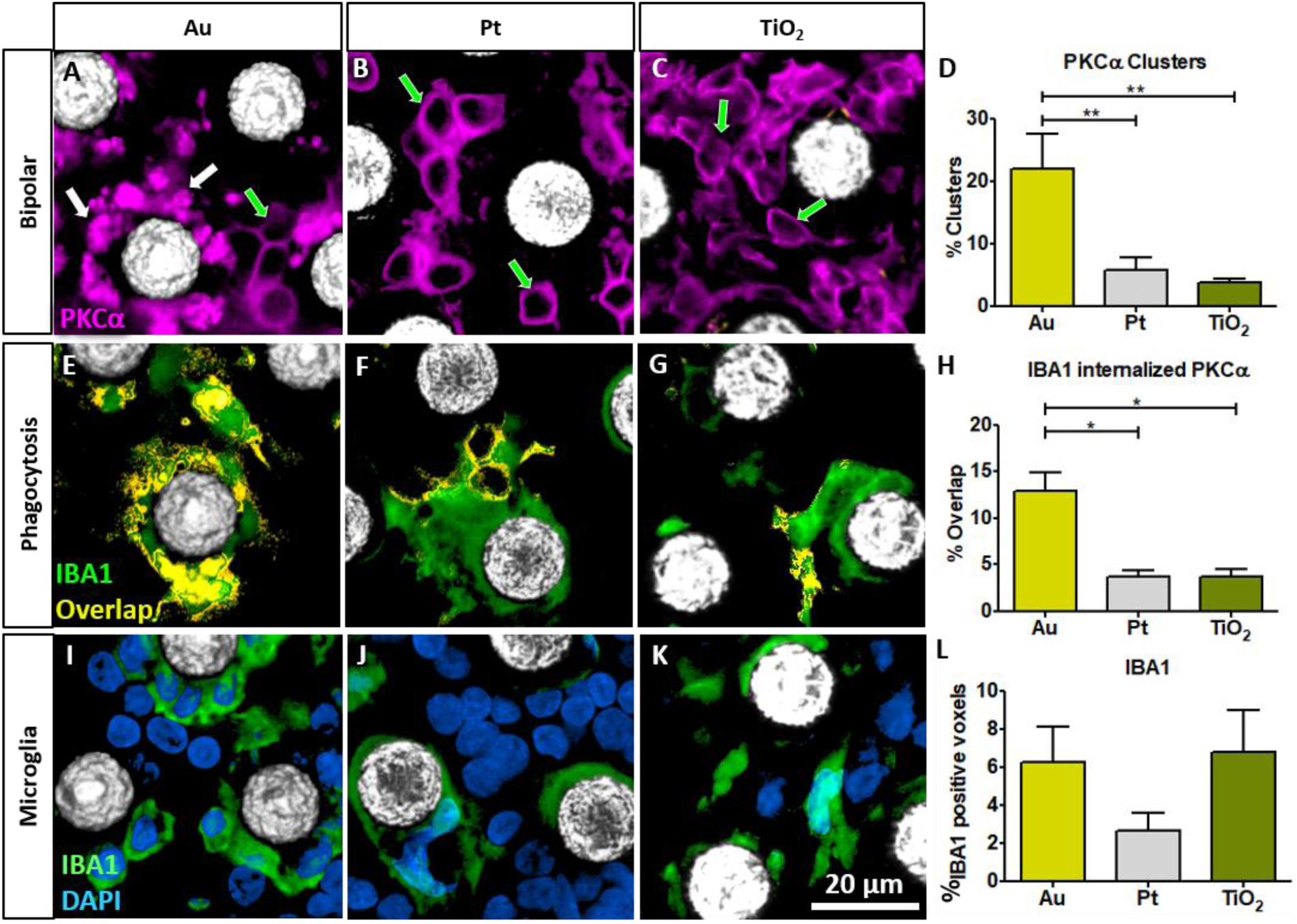
Representative confocal images of immunolabelled retinal wholemounts from RCS rats 6 weeks after subretinal implantation of pillar arrays coated with gold (Au; left column), platinum (Pt; middle column), or titanium dioxide (TiO₂; right column). The light reflection channel delineates the position of the implanted pillars from a top-down view (white). PKCα (magenta) labels rod bipolar cells. Green arrows indicate bipolar cells with preserved morphology, characterized by uniform cell membrane staining and a central nuclear void. White arrows highlight aberrant PKCα-positive clusters lacking normal bipolar cell morphology or nuclei. Such clusters are prominent in Au-coated implants (A) and largely absent in Pt (B) and TiO₂ (C) samples. D) Au-coated implants exhibit a significantly greater cluster area, expressed as a percentage of the total PKCα staining, compared with Pt and TiO₂ coatings, whereas no significant difference is detected between Pt and TiO₂. E-G) IBA1 (green) marks microglia, and its overlap with PKCα, shown in yellow, indicates internalization of bipolar cell material by microglia. Increased IBA1– PKCα overlap is evident in Au samples (E) relative to Pt (F) and TiO₂ (G), and is quantified in (H). In all groups (I-K), IBA1-positive cells accumulate around the pillars. (L) Total microglial presence, measured as the percentage of IBA1-positive voxels in the INL, shows no significant difference between different metals. Cluster - Au vs Pt: P = 0.0091 (**), Au vs TiO₂: P = 0.0054 (**), Pt vs TiO₂: (P =0.33) (ns); Phagocytosis - Au vs Pt: P = 0.021(*), Au vs TiO₂: P = 0.013 (*), Pt vs TiO₂: P = 0.47 (ns); Microglia - Au vs Pt: P = 0.33 (ns), Au vs TiO₂: P = 0.89 (ns), Pt vs TiO₂: P = 0.29 (ns).

In Au-coated implants, PKCα labelling revealed frequent aberrant clusters adjacent to pillar structures, characterized by aggregated PKCα-positive material lacking the typical bipolar cell membrane staining and nuclear void (Figure 7A). In contrast, Pt- and TiO₂-coated pillars demonstrated well-preserved bipolar cell morphology, with uniform membrane staining and intact somata surrounding the implants (Figures 7B–C). Quantification confirmed a marked increase in cluster area with Au compared to Pt and TiO₂ (Figure 7D: Au vs Pt: p = 1.9 × 10⁻⁶; Au vs TiO₂: p ≈ 6.6 × 10⁻⁸), whereas Pt and TiO₂ were similar (Pt vs TiO₂: P ≈ 1.0). These data indicate that Au-coated pillars induce structural disruption of rod bipolar cells. Given that bipolar cells are the target of stimulation with subretinal prosthesis, such morphological aberrations represent a potential compromise of the intended stimulation pathway.

To assess potential activation of the retinal immune system, IBA1 staining was applied to mark microglial cells. Overlap between IBA1 and PKCα staining reveals microglial internalization of bipolar cell-derived material. This overlap was significantly greater in Au (Figure 7E) compared with Pt and TiO₂ (Figure 7F-G, Au vs Pt: p = 8.02 × 10⁻⁶; Au vs TiO₂: p = 2.66 × 10⁻⁷), whereas Pt and TiO₂ again showed no difference (P = 0.938) (Figure 7H). The magnitude of IBA1–PKCα overlap closely mirrored the increase in PKCα-positive cluster formation, supporting the interpretation that microglia are engulfing bipolar cells and/or their remnants following exposure to Au. Gold may directly compromise bipolar cell integrity, generating acellular clusters that are subsequently recognized and cleared by microglia. Regardless the mechanism, increased microglial activation in the context of neuronal implants is undesirable. Activated microglia have been linked to bystander phagocytosis of stressed but viable neurons and to amplification of local inflammatory cascades [27, 28].

Despite clear differences in phagocytic activity, the total presence of microglia around pillars did not differ significantly among the three coatings (Figure 7I-L, Au vs Pt: p = 0.33; Au vs TiO₂: p = 0.89; Pt vs TiO₂: p = 0.29). IBA1-positive cells accumulated around all pillar types, suggesting that microglial recruitment to the implant site is a general response to the presence of 3D structures in the subretinal space, independent of the coating material. The difference therefore lies in the state and behavior rather than in the number of microglia, and Au uniquely perturbs the local cellular environment, shifting microglia from passive residency to active clearance of neuronal components.

### 2.5. Comparison of Pt and TiO_2_ coatings

Both TiO_2_ and Pt coatings demonstrated improved biocompatibility, as indicated by the reduced inflammatory response of the retina, compared to gold. However, Pt coating exhibited ten-fold higher capacitance than TiO_2_ (240 vs 25 gF/cm^2^), which means ten times higher current will flow from the sidewalls with Pt coating than with TiO_2_. The ratio of capacitance of the sidewalls, C_w_, to that of the SIROF-coated pillar tops, C_t_, depends on both the specific capacitance of each material and the corresponding surface areas, C_w_/C_t_ = (C_Pt_*A_w_)/(C_SIROF_*A_t_). For a cylindrical pillar with diameter D and height H, the sidewall area is A_w_ =π·D·H, while the top surface area is A_t_ =π·D^2^/4, so their ratio is A_w_/A_t_ = 4H/D. This geometric factor becomes particularly important for small pixels, where the pillar diameter is reduced and the relative sidewall area increases. For example, in 22 µm pixels with H = 40 µm and D = 8.5 µm, A_w_/A_t_ ∼19, in contrast, for 40 µm pixels the ratio is ∼10. When combined with the ratio of specific capacitances for Pt and SIROF (0.24 and 6 mF/cm², respectively), the sidewall capacitance becomes comparable to that of the pillar top, with the ratio C_w_/C_t_ reaching ∼0.75 and ∼0.4 for 22 µm and 40 µm pixels, respectively. To inject the charge predominantly from the pillar top, close to the INL, as intended [9], the specific capacitance of the sidewall should be decreased, and therefore, TiO_2_, with its 10-fold lower specific capacitance, is a better choice in this regard.

## 3. CONCLUSIONS

Although gold is a very convenient material for electroplating of microelectrodes, since retinal tissue exhibits an adverse response to gold, the pillar side walls should be insulated with biocompatible materials. Selective coating of the pillars side walls, while protecting the base of the implants, is quite challenging for conventional coating techniques, such as sputtering or atomic layer deposition.

We demonstrated that electrodeposition of Pt or TiO_2_ can selectively coat the pillar electrodes, while the return electrodes are protected by photoresist. Since the photosensitive areas separating the active and return electrodes are not conductive, protection of the return electrodes can be accomplished by noncritical photoresist masks, where the distance between the pillar and the edge of photoresist is not critical, as long as the pillar is exposed while the return electrode is covered. For the two approaches to patterning the photoresist, the etch-back process involves fewer processing steps, but requires careful optimization to expose the entire pillar sidewalls uniformly across the wafer, without compromising the protection of the return electrodes. An additional challenge can be the narrow openings between the pillars and photoresist restricting the access for the electroplating solution and leading to incomplete coating near the pillar base. In contrast, even though noncritical photolithography of a second photoresist requires several additional process steps, it provides better control of the coating pattern than the etch-back approach.

Both coatings demonstrated improved biocompatibility, but since TiO_2_ exhibited ten-fold lower capacitance than Pt, it is a better choice to ensure current injection predominantly from the SIROF-coated pillar tops. More broadly, the biocompatible coating techniques we developed for 3D electrodes are highly relevant not only for retinal prostheses, but for any implantable neural interfaces and bioelectronic devices that increasingly adopt three-dimensional architectures to improve proximity to target neurons. Both techniques, lithography and electrodeposition, can be used on a wafer scale and therefore offer versatile fabrication strategies for electro-neural interfaces of complex architecture on an industrial scale.

## 4. EXPERIMENTAL SECTION

### 4.1. Wafer Fabrication

The devices were fabricated on blank 4-inch Si/SiO₂ wafers without integrated diodes. Amorphous silicon (α-Si) was deposited by low-pressure chemical vapor deposition (LPCVD) for 90 min, and its resistivity was adjusted by PH3 doping using a flow rate of 2.5 SCCM (15% PH3 in SiH4). The α-Si resistors were first patterned using a standard photolithography technique, followed by the definition of Ti (50nm)/Au(200nm) active and return electrode structures, as previously reported [29]. A 230 nm-thick hydrogenated amorphous silicon carbide (a-SiC:H, referred to as SiC throughout the text) encapsulation layer was subsequently deposited by plasma-enhanced chemical vapor deposition (PECVD), the process parameters are outlined in ref. [30]. Vias were then opened through the SiC layer to expose the underlying active and return electrodes using inductively coupled plasma etching (ICP-RIE) performed in PlasmaTherm Metal Etcher (PT-MTL) system. Finally, a 300 nm thick platinum layer was deposited across the wafer to establish electrical connection for subsequent electroplating and electrodeposition steps.

Devices intended for animal implantation had additional 30 µm deep singulation trenches defined with deep reactive ion etching. The implants were released from the wafer by mechanical grinding followed by XeF_2_ etching. To encapsulate exposed Si, the back side of the implants was coated with a 200 nm sputtered layer of material of interest for biocompatibility experiments, namely Pt for the Pt group, Au for the Au group and Ti for the TiO_2_ group.

### 4.2. Fabrication of 3D Gold Pillar Electrodes

High-aspect ratio gold electrodes were fabricated by electroplating following a previously reported process. A 30 µm thick negative photoresist (KMPR-1025) was patterned by contact photolithography (Karl Suss MA6) to define the electroplating mask followed by the development in a TMAH-based developer. To ensure exposure of the underlying seed electrode, a descum step was performed using a reactive ion etching tool. Next, the patterned wafers were mounted in a custom PTFE wafer holder designed to isolate the wafer edges and back surface. Electrical contact was established via a conductive handle integrated into the holder, while a platinized titanium mesh positioned parallel to the wafer surface served as the anode. Gold deposition was carried out in a commercial electroplating solution (NB Semiplate AU 100TH, NB Technologies, Bremen, Germany) at a temperature of 30 °C and constant agitation at 40 rpm. A constant current density of 1 mA cm^-2^ was applied throughout the process, resulting in a deposition rate of approximately 3 µm per hour. Once the desired gold structure height was achieved, the wafers were removed from the plating solution and rinsed thoroughly with deionized water. The KMPR-1025 resist mask was then removed by immersion in PG remover at 80 °C, leaving free-standing three-dimensional gold pillar structures.

### 4.3. Etch back process of KMPR

The PR1 mask used for electroplating (KMPR-1025) can be etched away using an RIE process including oxygen and fluorine. An etch was developed for PR1 removal with a recipe of 60sccm SF6, 10sccm O2 at 150W with a pressure of 0.1 Torr. By controlling the etch time, a process was developed where in photoresist remains above the bulk of the wafer, but is removed surrounding each electroplated gold pillar. This developed process results in a new electrodeposition mask which selectively exposes three-dimensional structures, allowing for further processing.

### 4.4. Noncritical photolithography mask

The PR2 mask was created by maskless laser lithography: First, the negative photoresist was spray-coated to form a uniform film with the thickness of ∼ 6 µm. Next, the chip was exposed in regions between the pillars designed to protect the return electrodes while selectively opening the pillar sidewalls for subsequent electrodeposition processing. Patterning was performed using an ML3 MicroWriter system with AZ 15nXT (450cps) negative photoresist.

### 4.5. Electrodeposition of TiO2 coating

Before TiO_2_ deposition, the Au electrodes were cleaned using O_2_ plasma treatment. TiO_2_ films were grown on Au by anodic oxidative hydrolysis of an aqueous 50 mM TiCl_3_ solution adjusted with NaOH to pH ∼ 2.30 ± 0.05. The Au surface was held at 0 V vs Ag/AgCl reference electrode, corresponding to an increased bias offset of ∼ 450mV relative to the open circuit potential recorded prior to deposition (see Supplementary Information for more details) [20, 22]. The process was performed under nitrogen environment to avoid oxidation. The solution was continuously mixed to ensure uniformity. The growth process was controlled by time. See Supplementary Note S8 for more details.

### 4.6. Electrodeposition of Pt coating

After a mask for further processing has been utilized, be it etch back of PR1 or a noncritical photolithography mask, platinum electrodeposition was carried out to form a biocompatible coating. Similarly, to the deposition of TiO_2_, an O_2_ plasma treatment was used to clean the surfaces to be coated. Platinum was electrodeposited in a two-electrode set-up using chloroplatinic acid, mixed at a ratio of 1% chloroplatinic acid, 0.08% lead acetate and 98.92% DI water. The solution was constantly agitated with a PTFE stirrer and heated to a temperature of 70 °C. The Au pillar electrodes are the working electrodes and a platinized titanium mesh was used as the counter electrode. A current density of 1mA/cm^2^ was applied to grow the platinum layer, based on the exposed pillar surface area. The potential of the working electrode settles at approximately -100mV with respect to the platinized titanium mesh counter electrode. Electroplating was carried out for 10 minutes, resulting in a layer of thickness ∼18-19 nm as confirmed with atomic force microscopy.

### 4.7. Electrochemical characterization

All cyclic voltammetry (CV) scans were performed in phosphate buffered saline (PBS) solution in a three-electrode configuration using a large-area platinum electrode, an Ag/AgCl reference electrode and fabricated samples set as the working electrode. All measured potentials are reported with respect to the Ag/AgCl electrode. The working electrode potential sweep was between -0.6 and 0.6 V with 100 mV s*^-^*^1^ scan rate consistent with previously reported studies [25]. The selected potential range was chosen to reflect the maximum photovoltage typically achieved during the standard operation of the retinal implant. Furthermore, the selected range avoids Faradaic reactions including water reduction, hydrogen and oxygen evolution reactions [13, 25, 31].

### 4.8. Capacitance analysis

Double-layer capacitance C*di* was determined from cyclic voltammetry measurements acquired in a non-Faradaic potential window, where the anodic and cathodic currents appear parallel to each other and symmetrical around the zero current [25]: 0.1 – 0.3V for electrodeposited TiO_2_, 0.2 – 0.3V for electrodeposited and sputtered Pt, and -0.1V – 0.05V for evaporated Au, as described earlier [25]. The double-layer capacitance *(C dl*) was calculated using the following: where *l_an_* and *I_cat_* are the average values of anodic and cathodic currents, respectively. The scan rate of all CV measurements, *v,* was set to 100 mV s^-1^. See Supplementary Note S9 for representative CV curves of the analyzed materials.

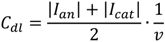

### 4.9. Animals and Surgical Procedures

All animal experiments were approved by the Stanford Administrative Panel on Laboratory Animal Care (APLAC) and were performed in accordance with institutional guidelines and the Association for Research in Vision and Ophthalmology (ARVO) Statement for the Use of Animals in Ophthalmic and Vision Research. Experiments were conducted using rats from a Royal College of Surgeons (RCS) colony maintained at the Stanford animal facility. For quantitative analysis, each coating group (Au, Pt, and TiO₂) contained a minimum of 12 samples. Implantations were performed after postnatal day 180 (P180), when photoreceptor degeneration in this model is complete. Animals were anesthetized by intraperitoneal injection of ketamine (75 mg/kg) and xylazine (5 mg/kg). A 1.5-mm incision was created through the sclera and choroid approximately 1 mm posterior to the limbus. Retinal detachment was induced by injection of sterile saline into the subretinal space, after which the implant was inserted into the created bleb at least 3 mm from the entry site. The conjunctiva was closed using 10-0 nylon sutures, and topical antibiotic ointment (bacitracin/polymyxin B) was applied postoperatively. Proper implant placement and retinal reattachment were verified using optical coherence tomography (Spectralis HRA2-OCT, Heidelberg Engineering, Heidelberg, Germany).

### 4.10. Retinal Immunohistochemistry

Animals were euthanized 6–9 weeks after implantation by intracardiac injection of phenytoin/pentobarbital (VetOne Euthanasia Solution). Eyes were enucleated and rinsed in phosphate-buffered saline (PBS). The anterior segment and lens were removed, and a 3 × 3 mm retinal region centered on the implant was isolated and fixed in 4% paraformaldehyde (Electron Microscopy Sciences, PA, USA) for 12 h at 4 °C. Tissue samples were permeabilized in 1% Triton X-100 (Sigma-Aldrich, CA, USA) in PBS for 3 h at room temperature, followed by blocking in 10% bovine serum albumin (BSA) for 1 h. Primary antibodies against PKCα (Thermo Fisher Scientific) and IBA1 (Wako, Japan) were applied for 48 h at room temperature in a solution containing 0.5% Triton X-100 and 5% BSA in PBS. After washing for 6 h in PBS containing 0.1% Tween-20 (PBS-T), samples were incubated for 48 h at room temperature with secondary antibodies (donkey anti-mouse Cy3, Jackson ImmunoResearch; donkey anti-rabbit Alexa Fluor 488, Thermo Fisher Scientific). Nuclei were counterstained with DAPI. Samples were washed in PBS-T for 4–6 h and mounted using Vectashield mounting medium (Vector Laboratories, Burlingame, CA, USA).

### 4.11. Whole-Mount Retinal Imaging and Image Processing

Three-dimensional imaging of retinal whole mounts was performed using a Zeiss LSM 880 inverted confocal microscope with ZEN Black acquisition software. Implant structures were visualized using the reflection signal generated by a 514-nm laser, detected through a neutral-density beam splitter configured for 80% transmission and 20% reflection. Z-stack images were acquired through the full retinal thickness, with the upper boundary defined at the inner limiting membrane and the lower boundary extending 10 µm below the base of the implant structures. Image stacks were collected from the central region of each quadrant using a 40× oil-immersion objective with acquisition fields exceeding 225 × 225 µm and z-step intervals of 380–470 nm. Signal attenuation within implant structures was compensated using the Z-stack correction module in the ZEN software. Confocal image stacks were processed using the Fiji distribution of ImageJ. To compensate for intensity variations along the z-axis, contrast in each XY plane was adjusted to achieve 0.3% saturation. Images were filtered using a median despeckling operation, and background fluorescence was reduced using a rolling-ball subtraction algorithm. Additional gamma and min–max adjustments were applied where necessary to further suppress background noise. Gaussian smoothing was applied to nuclear staining channels to reduce intra-cellular intensity fluctuations. Implant geometry was reconstructed by extending the reflection signal toward the extraocular side of the stack.

### 4.12. Quantification of immunohistochemistry

Three-dimensional segmentation of implant structures was performed based on the reflection channel using the Moore–Neighbor tracing algorithm implemented in the MATLAB “bwboundaries” function (MATLAB R2021b, MathWorks). Control image stacks without implants were analyzed as a single segment. For IBA 1 quantification, voxels with intensities exceeding 15% of the maximum signal were classified as positive. PKCα clusters, defined as signal forming an acellular structure lacking a nucleus in the center, were manually segmented to distinguish from signal corresponding to health bipolar cells. The total cluster areas at 2 heights in INL of each samples were calculated, averaged and plotted as a percentage of the total PKCα signal. Overlap of the IBA1 signal with PKCα was used to indicate the macrophages engulfing/phagocytosing bipolar cells. The overlap signal was isolated by using the image calculator ‘AND’ function in imageJ to create a new channel with only corresponding overlap signal it and the percentage overlap was calculated as described for the PKCα signal.

## Acknowledgments

Studies were supported by the National Institutes of Health (R01-EY-035227 and P30-EY-026877), Department of Defense (W81XWH-22-1-0933) and AFOSR (FA9550-24-1-0138). Part of this work was performed at the Stanford Nanofabrication Facility (SNF) and the Stanford Nano Shared Facilities (SNSF), supported by the National Science Foundation under award ECCS-2026822. KM was supported by an RAEng Chair in Emerging Technologies. EB was partially supported by the Rhona Reid Charitable Trust.

## Conflicts of Interest

The authors declare no conflicts of interest.

## Notes

### Competing Interest Statement

The authors have declared no competing interest.

### Summary of Updates

The revised version provides embedded figures for improved readability.

