## Supplementary_Information for "Electrodeposited Biocompatible Coatings for 3D Electrodes on Retinal Prostheses"

**Supplementary Note S1: Finite-element modeling of electric potential above the photovoltaic arrays**

To evaluate electric potential generated in the electrolyte above the photovoltaic arrays, a finite-element analysis model was developed using COMSOL Multiphysics 5.6. This model uses a circuit model to define the photocurrent generated by a train of 4 ms pulses of NIR light (880 nm wavelength) at 4 $mW/mm^2$ irradiance. An electrolyte of 1000 Ohm*cm resistivity is assumed as a model of retinal tissue. The electrode-electrolyte interface is a linearized version of the Butler-Volmer law, with the double layer capacitance values and exchange current density defined for each electrode material. For the Pt-coated sidewalls of 30 μm tall pillar electrodes, $C_{Pt}$ = 100 μF $cm^{-2}$, whilst for the 400 nm thick layer of SIROF on top of the pillars, $C_{SIROF}$= 8.52 mF $cm^{-2}$. The honeycomb mesh of return electrodes is also modelled as a 400 nm-thick and 4 μm wide SIROF layer. All other surfaces which make up the modelled device are set as insulating boundaries.

The color map in Figure S1 shows the electric potential throughout the modelled retinal tissue domain. Full details of this model are found in Butt, E., et al., *Three-dimensional electro-neural interfaces electroplated on subretinal prostheses*. J Neural Eng, 2024. **21**(1).

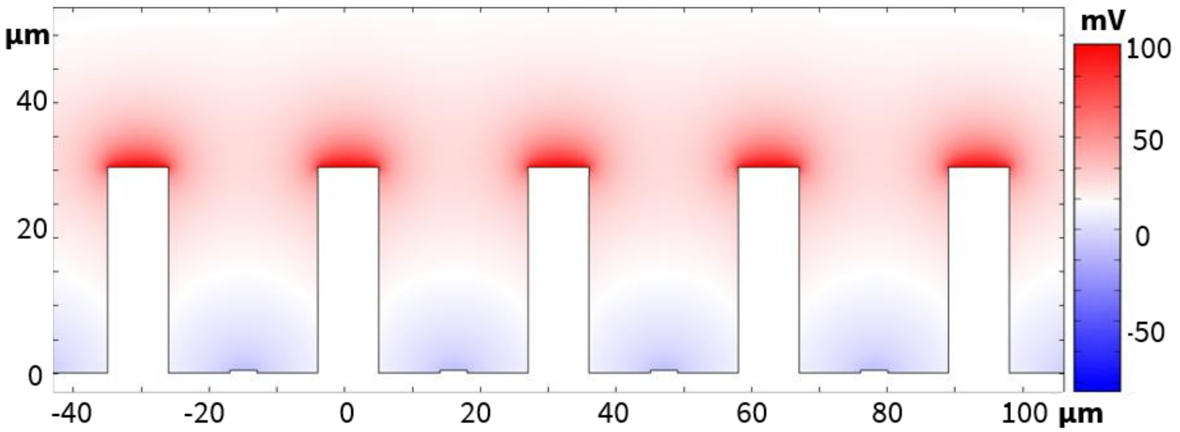


**Supplementary Figure S1.** Electric potential in electrolyte above the photovoltaic array with 30μm pixels and 30μm tall pillars on active electrodes. Color map indicates the electric potential with respect to axonal terminals of bipolar cells at 90 μm height - in the middle of the inner plexiform layer.

**Supplementary Note S2: Elemental analysis of $TiO_2$ films**

To establish the best conditions for observation of $TiO_2$ with energy-dispersive X-ray spectroscopy, EDS measurements were performed using different accelerating voltages, including 5,10 and 20 kV. An accelerating voltage of 10 kV was selected for subsequent measurements because it provides clearly resolved Ti and O features while minimizing the contribution from the underlying Si substrate signal (Supplementary Figure S2.1. A,B). EDS mapping was performed across the uncoated gold (Au) and $TiO_2$-coated ($TiO_2$/Au) regions. The Ti and O maps show a clear increase in signal within the $TiO_2$-coated region, consistent with the localized growth of the electrodeposited $TiO_2$ film. In contrast, the Au signal is reduced in the coated area because the $TiO_2$ layer attenuates the EDS signal originating from the underlying Au electrode. Similar EDS measurements were also performed on the $TiO_2$-coated pillars, confirming the presence of Ti and O - the elemental composition of the electrodeposited $TiO_2$ coating on sidewalls.

Composition of the electrodeposited $TiO_2$ films was further evaluated by X-ray photoelectron spectroscopy (XPS) combined with sputtering for bulk signal access. XPS measurements were performed using a monochromatic Al Kα source operated at 46.1 W with a 200 μm spot size and a 45° photoelectron take-off angle. XPS typically probes only the upper ~5–10 nm of a film, but the surface typically has higher concentration of oxygen and contains adsorbed carbon species or other process residues. Therefore, depth profiling and surface cleaning were carried out with a gas cluster ion beam (GCIB) at 5 kV (2500-atom clusters). Supplementary Figure S2.2. presents the sputter series collected at the film surface before and after iterative sputtering. Besides the survey spectra, high-resolution spectra of Ti 2p and O 1s were obtained as presented in the main text. After 3 min of GCIB sputtering, quantification from the high-resolution scans reveals that the $TiO_2$ films exhibited a composition of approximately 67 at.% O and 33 at.% Ti, consistent with stoichiometric $TiO_2$.

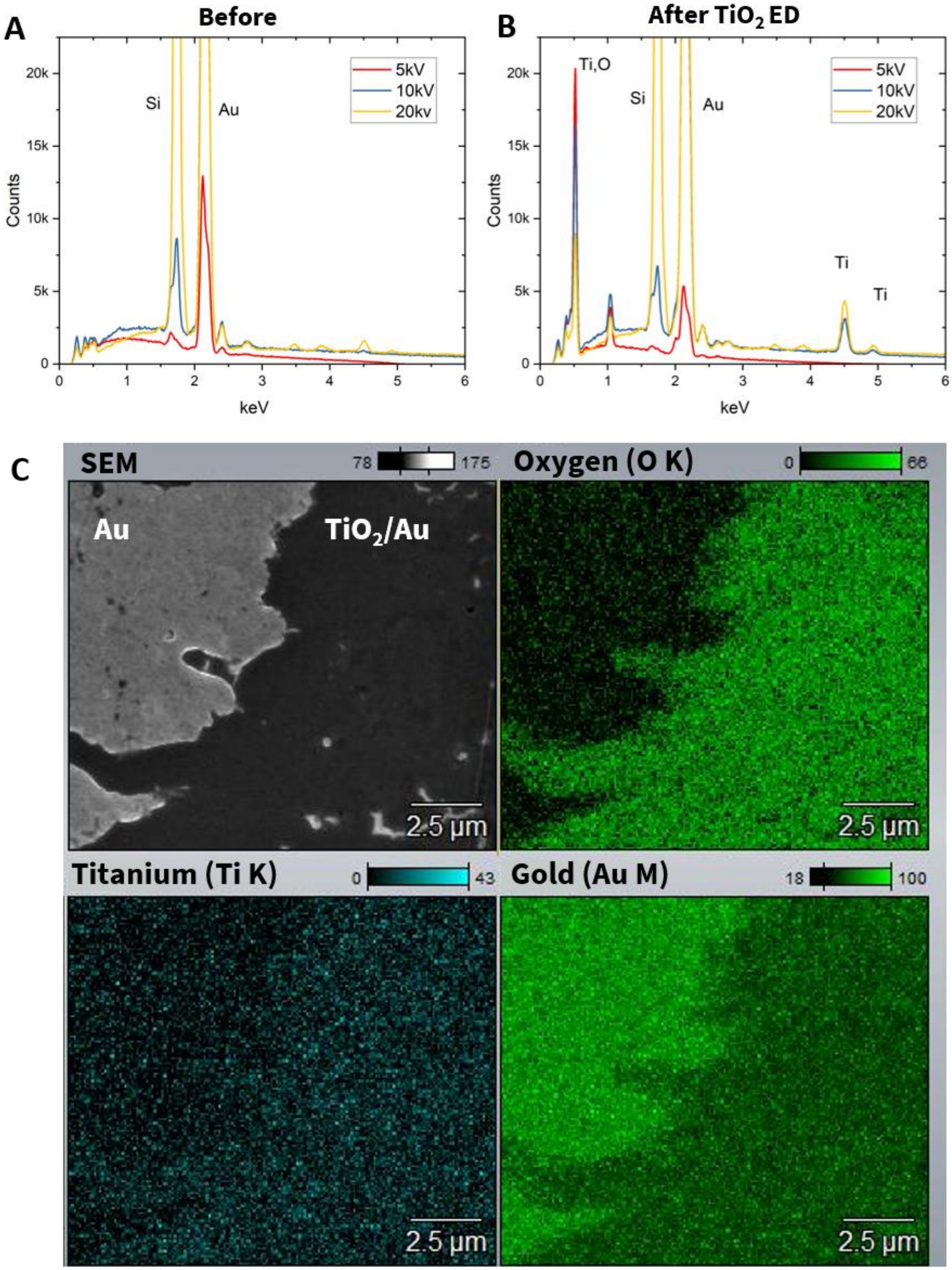


**Supplementary Figure S2.1.** EDS characterization of electrodeposited $TiO_2$ films. (A,B) EDS spectra acquired at accelerating voltages of 5, 10 and 20 kV before and after $TiO_2$

electrodeposition, respectively. The 10 kV condition provides clear Ti and O signals while limiting the contribution from the Si substrate. (C) SEM image and corresponding EDS elemental maps (O, Ti and Au) acquired across the boundary between the $TiO_2$-coated and uncoated Au regions. The Ti and O signals are localized within the $TiO_2$-coated region, while the Au signal is correspondingly attenuated. Note that the boundary in this sample was defined using Kapton tape resulting in a relatively rough edge.

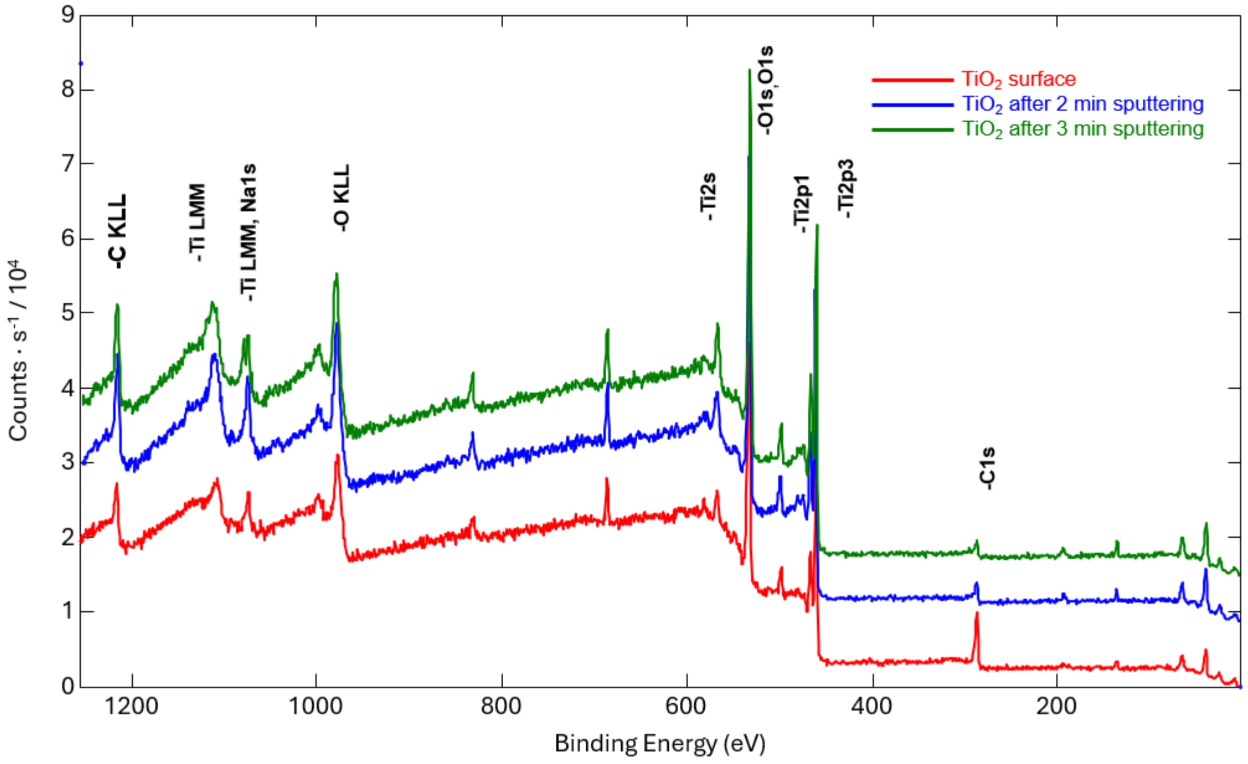


**Supplementary Figure S2.2.** XPS spectra of electrodeposited $TiO_2$ films before and after GCIB sputtering, which removes the surface layer, enabling evaluation of the film composition in the bulk.

**Supplementary Note S3: Effect of electrodeposition time on the cyclic voltammetry measurements of $TiO_2$ films**

To evaluate the effect of electrodeposition time on the electrochemical properties of the $TiO_2$ coatings, cyclic voltammetry measurements (CV scans) were performed on films electrodeposited for 5, 15, 20, 30 and 60 min. All $TiO_2$ coatings showed significantly lower current densities than the uncoated Au electrode, consistent with the insulating properties of high band gap semiconductor $TiO_2$. Interestingly, even after 5 mins of $TiO_2$ deposition the characteristic electrochemical peaks of Au were fully suppressed and current density at -0.6V drastically dropped. Only small differences were observed among longer coated $TiO_2$ films indicating that electrodeposition time did not substantially change the electrochemical properties of the films. Similarly, surface topography investigated with AFM and discussed in the main text showed no change in the RMS roughness with increasing deposition time.

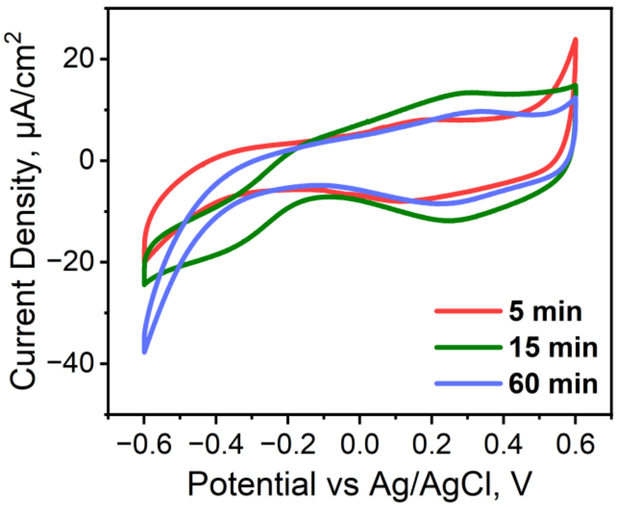


**Supplementary Figure S3.** Cyclic voltametry curves for a bare Au substrate and for $TiO_2$ films electrodeposited for 5, 15 and 60 min. All electrodeposited $TiO_2$ films show suppression of characteristic Au peaks and reduced current density, with only minor variations among films with different deposition times.

**Supplementary Note S4: Effect of the $TiO_2$ staining on reflectance of the photosensitive region.**

To check whether the thin $TiO_2$ staining layer observed at the pillar base (Figure 3F in the main text) will affect the device performance, we measured changes in reflectance of the photosensitive diode region. Chips from different wafer locations were exposed to the $TiO_2$ electrodeposition solution under conditions identical to those used for coating of the pillar devices. For each chip, one half was protected with photoresist (PR), while the other half was directly exposed to the electrodeposition solution. Following exposure, the chips were rinsed with DI water, the photoresist was removed with acetone, and the samples were subsequently rinsed with IPA and dried under $N_2$ gun flow. Reflectance spectra were collected before and after exposure, comparing the directly exposed and PR-protected regions of each chip. No significant change in reflectance was observed within the electrodeposition time range used to obtain the desired coating thickness (15 min). The small reflectance differences between chips originate from wafer-scale variation in the thickness of the SiC layer.

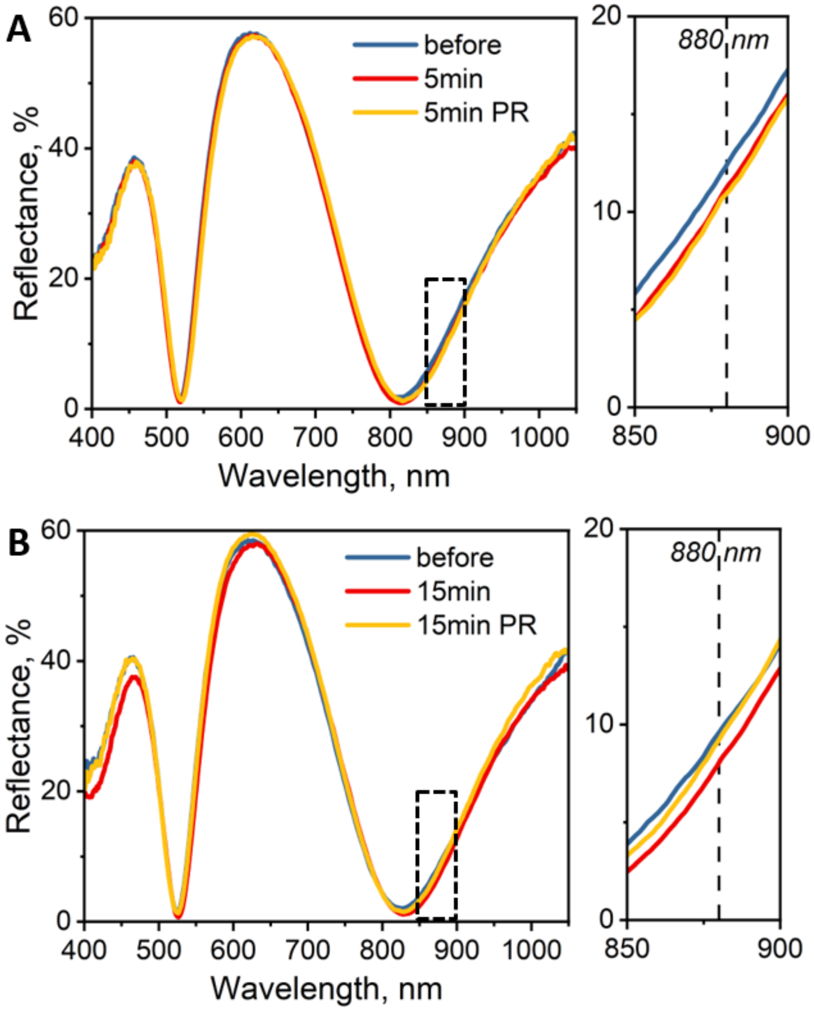


**Supplementary Figure S4.** Reflectance spectra for the device regions passively exposed to the $TiO_2$ electrodeposition solution for 5 min (A) and for 15 min (B). The areas protected with photoresist (PR) are used as a control of the reflectance. Zoomed-in regions are displayed for analysis of reflectance changes at stimulation wavelength 880 nm. The chips from different parts of the wafer showcase the typical variation in reflectance across the wafer due to SiC thickness variation.

**Supplementary Note S5: Surface roughness of Pt films made by pulsed electrodeposition**

To reduce the roughness of electrodeposited platinum films, we explored using the pulsed electrodeposition (2.5Hz pulse frequency, 50% duty cycle and 3mA $cm^{-2}$ current density). However, the atomic force microscopy (AFM) analysis showed no improvement in film roughness compared with films deposited under continuous conditions. An average root-mean-square (RMS) roughness of ~7 nm was observed under both electrodeposition conditions.

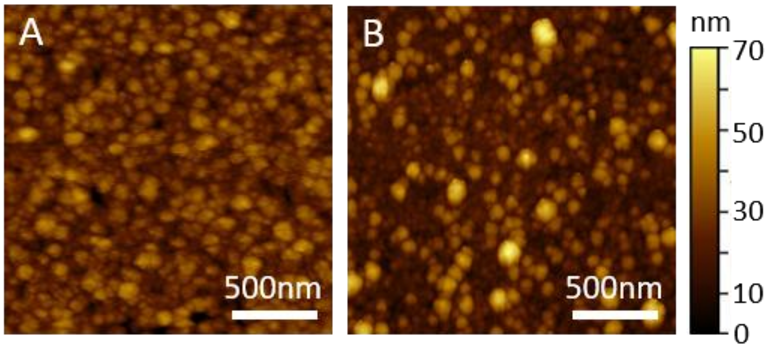


**Supplementary Figure S5.** AFM maps for electrodeposited Pt films on smooth Au-coated substrates. The Pt films were prepared using pulsed (A) and continuous (B) deposition settings. No improvement in roughness was observed for pulsed deposition, as measured by RMS value, reporting ~7 nm for both conditions.

**Supplementary Note S6: Elemental analysis of electrodeposited Pt films**

Similarly to the electrodeposited $TiO_2$ films, we used XPS to characterize the elemental composition of the electrodeposited Pt films. The XPS acquisition and sputtering conditions were identical to those used for $TiO_2$ films, described in Supplementary Note S2. Besides the survey spectra, high-resolution spectra of Pt and O 1s were obtained, as presented in the main text. Stoichiometric analysis following 6 min of sputtering showed that the bulk film consisted predominantly of metallic Pt (96.5 at.% with only 3.5 at.% O).

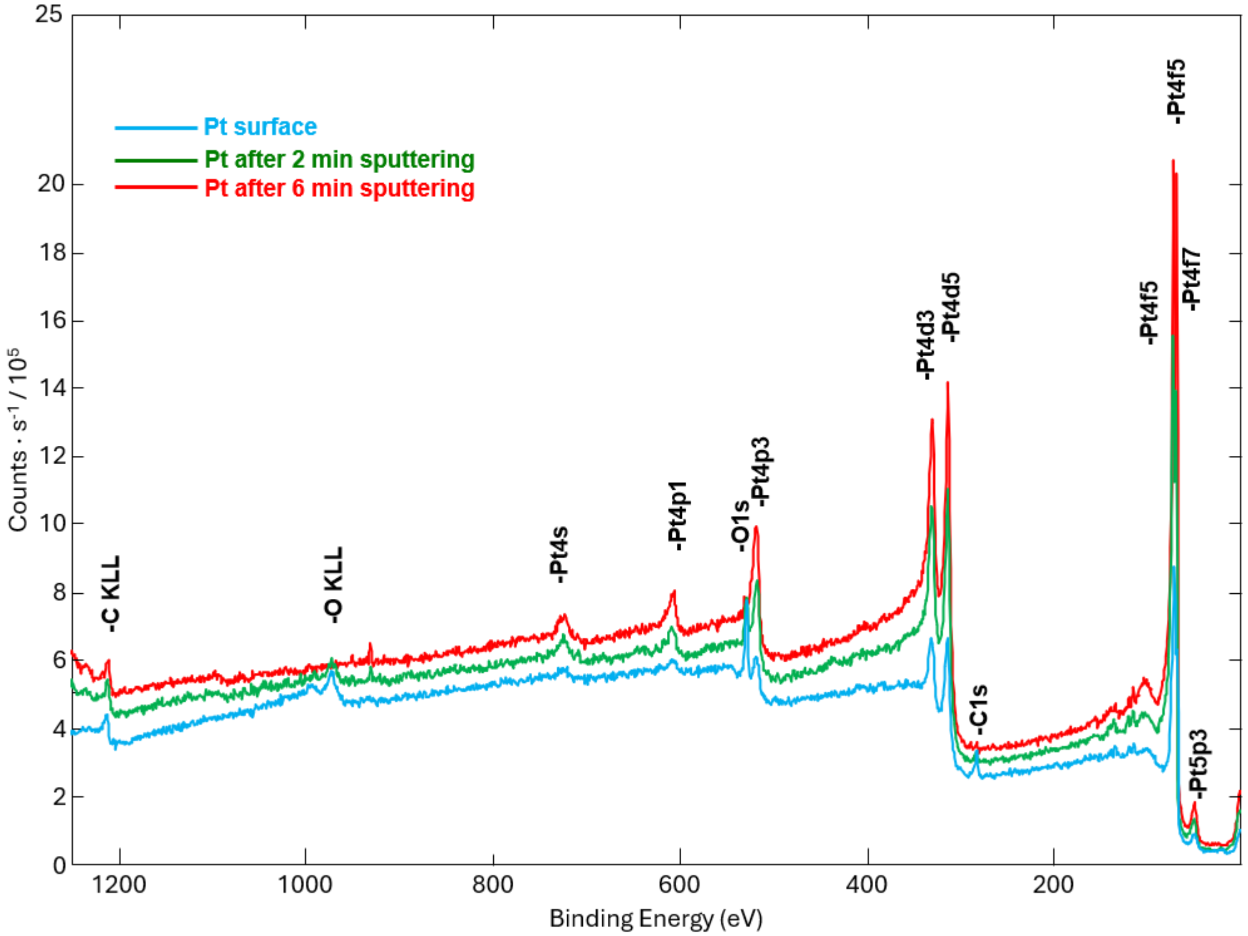


**Supplementary Figure S6.** XPS spectra of electrodeposited Pt films before and after GCIB sputtering for 2 and 6 mins. Sputtering removes the surface layer, enabling evaluation of the bulk film composition.

**Supplementary Note S7: Continuity of electrodeposited Pt coatings along the pillar sidewalls.**

To assess the continuity of the electrodeposited Pt coating along the pillar sidewalls, we performed spatially correlated SEM-EDS measurements at multiple positions on the pillars. Particular attention was given to the pillar base, where access of the electrodeposition solution to the Au surface is most restricted. SEM imaging was used to evaluate the morphological continuity of the coating by confirming the absence of visible cracks or gaps, which could expose Au regions. Corresponding EDS analysis verified that the elemental composition of the coating is Pt. Because XPS averages the elemental composition over a relatively large area (~100 μm), we used EDS to evaluate the coating composition locally at the microscale. We first established the EDS signature of electrodeposited Pt using planar samples, as discussed in the main text. Uncoated Au exhibits a dominant Au Lα peak at 9.7 keV, whereas Pt-coated Au shows an additional Pt Lα peak at 9.4 keV. This Pt-specific feature was then used to evaluate the presence of the Pt coating at different locations along the pillar sidewalls, including the pillar base, as shown in the Figure below. Together, the SEM and EDS measurements indicate that a continuous Pt coating extends along the pillar sidewalls down to the base.

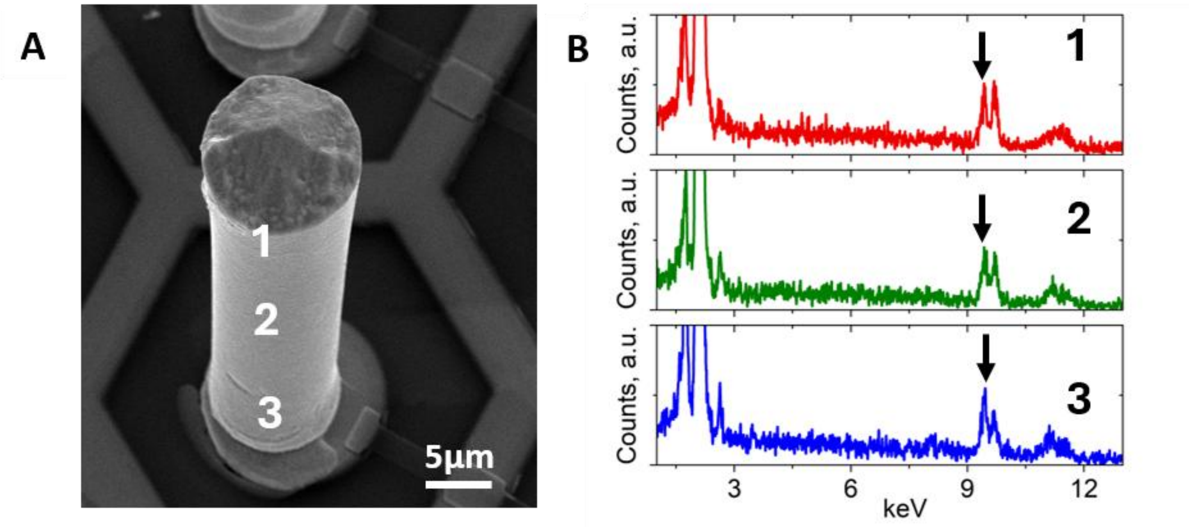


**Supplementary Figure S7.** Correlative SEM-EDS analysis of electrodeposited Pt coatings. EDS spectra are collected at different positions along the pillar sidewall. The Pt signature peak at 9.4 keV, highlighted by the arrows, confirms presence of the Pt coating along the pillar.

**Supplementary Note S8: Electrodeposition process for $TiO_2$ films**

Titanium trichloride ($TiCl_3$) and sodium hydroxide (NaOH) were obtained from Sigma-Aldrich. All chemicals were used as received. Deionized (DI) water (>18 MΩ·cm) was used in all experiments. The process starts by preparing 50mM solution of $TiCl_3$ adjusted with NaOH to reach pH of ~2.3±0.5. The acidic solution is prepared under a nitrogen atmosphere and constant mixing. First, DI water is added to achieve the 50mM concentration, followed by the slow introduction of NaOH with continuous monitoring of the solution pH. After the solution reaches the target pH, it is transferred into the electrodeposition cell for film growth. The final solution appears dark blue. If the solution is left under ambient environment, it oxidizes and changes color.

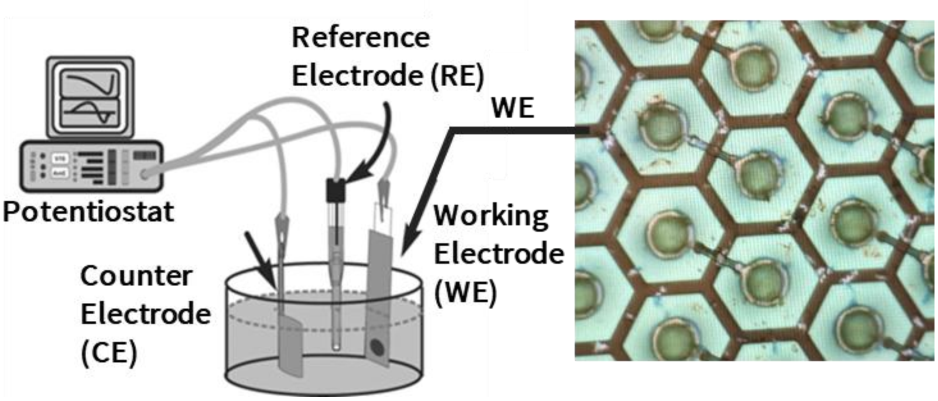


**Supplementary Figure S8.** Schematic of the three-electrode potentiostat setup used for electrodeposition of $TiO_2$ films.

**Supplementary Note S9: Measurement of the double-layer capacitance**

The double-layer capacitance of the investigated materials was determined from cyclic voltammetry (CV) measurements performed within a narrow potential window dominated by non-faradaic charging (see the main text for details). All CV measurements were performed at a scan rate of 100 mV $s^{-1}$. Representative CV curves and the corresponding extracted capacitance values are shown in the Figure below.

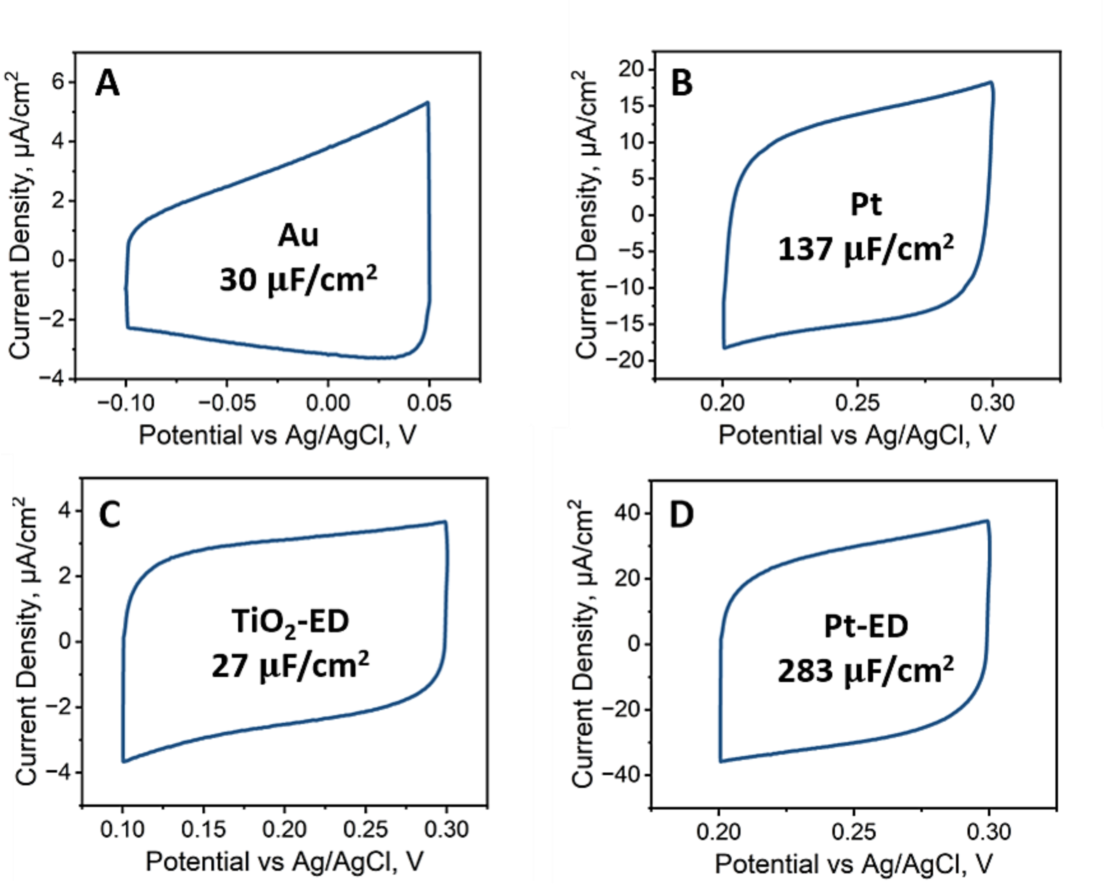


**Supplementary Figure S9.** Cyclic voltametry curves used to estimate the double-layer capacitance of (A) electron-beam evaporated Au, (B) sputtered Pt, electrodeposited $TiO_2$ (C) and electrodeposited Pt (D). Electrodeposited $TiO_2$ exhibited the lowest capacitance, while electrodeposited Pt showed an increased capacitance relative to the sputtered reference Pt film, as discussed in the main text.
